# Medial frontal theta is entrained to rewarded actions

**DOI:** 10.1101/144550

**Authors:** Linda M. Amarante, Marcelo S. Caetano, Mark Laubach

## Abstract

Rodents lick to consume fluids. The reward value of ingested fluids is likely to be encoded by neuronal activity entrained to the lick cycle. Here, we investigated relationships between licking and reward signaling by the medial frontal cortex [MFC], a key cortical region for reward-guided learning and decision-making. Multi-electrode recordings of spike activity and field potentials were made in male rats as they performed an incentive contrast licking task. Rats received access to higher and lower value sucrose rewards over alternating 30 sec periods. They learned to lick persistently when higher value rewards were available and to suppress licking when lower value rewards were available. Spectral analysis of spikes and fields revealed evidence for reward value being encoded by the strength of phase-locking of a 6-12 Hz theta rhythm to the rats’ lick cycle. Recordings during the initial acquisition of the task found that the strength of phase-locking to the lick cycle was strengthened with experience. A modification of the task, with a temporal gap of 2 sec added between reward deliveries, found that the rhythmic signals persisted during periods of dry licking, a finding that suggests the MFC encodes either the value of the currently available reward or the vigor with which rats act to consume it. Finally, we found that reversible inactivations of the MFC in the opposite hemisphere eliminated the encoding of reward information. Together, our findings establish that a 6-12 Hz theta rhythm, generated by the rodent medial frontal cortex, is synchronized to rewarded actions.

**Significance Statement:** The cellular and behavioral mechanisms of reward signaling by the medial frontal cortex [MFC] have not been resolved. We report evidence for a 6-12 Hz theta rhythm that is generated by the MFC and synchronized with ongoing consummatory actions. Previous studies of MFC reward signaling have inferred value coding upon temporally sustained activity during the period of reward consumption. Our findings suggest that MFC activity is temporally sustained due to the consumption of the rewarding fluids, and not necessarily the abstract properties of the rewarding fluid. Two other major findings were that the MFC reward signals persist beyond the period of fluid delivery and are generated by neurons within the MFC.

## Introduction

Reward-related neuronal activity is commonly found in the medial frontal cortex [MFC], aka mPFC, of humans (Glascher et al., 2009; Levy and Glimcher, 2011), primates (Watanabe, 1996; Shidara and Richmond, 2002; Roesch and Olson, 2004; Amiez et al., 2006; Padoa-Schiappa and Assad, 2006; Hayden et al., 2009; Luk and Wallis, 2009; Bouret and Richmond, 2010; Cai and Padoa-Schioppa, 2012), and rodents (Petyko et al., 2009; Horst and Laubach, 2012, 2013; Donnelly et al., 2014; Petyko et al., 2015). However, the behavioral determinants of these signals are not understood. In neurophysiological studies in experimental animals, rewards are typically given as liquids (Apicella et al., 1991; Carelli and Deadwyler, 1994) to avoid issues with chewing and grinding foods. A fixed cycle of specific behaviors (jaw opening, tongue protrusion and retraction, jaw closing, swallowing) underlies the processing of liquid rewards. These behaviors should have a major impact on reward signaling by the MFC.

Indeed, Horst and Laubach (2012) reported that MFC activity is sharply modulated when thirsty rats lick to consume water rewards in a MFC-dependent working memory task (Horst and Laubach, 2009). By modifying the task to delay the delivery of water on some trials, a subpopulation of MFC neurons was revealed that were selectively activated by the initiation of licking (Horst and Laubach, 2013). These changes in spike activity are accompanied by prominent fluctuations of MFC local field potentials [LFPs], specifically near the rats’ licking frequency (5-7 Hz). These signals were most prevalent in the most rostral MFC. A subsequent study found that reversible inactivation of this rostral part of the MFC dramatically reduces the duration of licking bouts (Parent et al., 2015a), similar to how inactivation of the more caudal MFC leads to excessive premature responding in tasks that require actions to be sustained over delay periods (e.g., Narayanan et al., 2006). These studies suggest that the rostral MFC is specialized for the value-guided control of consummatory behavior.

The goal of the present study was to determine if licking-related neuronal activity in the rostral MFC is sensitive to the reward value of ingested fluids. To examine this issue, we used a simple take-it-or-leave decision-making task, called the Shifting Values Licking Task (Parent et al., 2015a,b), to study reward signaling in relation to ongoing consummatory actions. Rats lick on a spout to receive liquid sucrose rewards and the concentration of sucrose alternates between a higher (better) and lower (worse) option every 30 sec. After only a few days of training, the rats learn to persistently lick when the better option is available and to suppress licking when the worse option is available. Bilateral reversible inactivations of the rostral MFC impair performance in this task (Parent et al., 2015a), resulting in temporally fragmented licking (dramatic reductions in the duration of licking bouts). Opposite effects are found after intra-MFC infusions of drugs that are known to enhance neuronal excitability near the licking frequency, such as the M-channel blocker XE-991 (Hu et al., 2002), and the “hunger hormone” ghrelin (Parent et al., 2015b).

To examine how the rostral MFC encodes reward information and controls value-guided consumption, we recorded spike activity and local field potentials [LFP] as rats performed the Shifting Values Licking Task. We found that neuronal activity in the MFC is entrained to the lick cycle when animals consume liquid sucrose rewards. These signals develop with experience, persist beyond periods of fluid delivery, and depend on MFC neurons. Together, our findings suggest that a 6-12 Hz rhythm generated by MFC neurons tracks engagement in reward-based consummatory behavior and encodes reward information.

## Materials and Methods

Procedures were approved by the Animal Use and Care Committees at the John B. Pierce Laboratory (where some of the experiments were conducted) and American University and conformed to the standards of the National Institutes of Health Guide for the Care and Use of Laboratory Animals. All efforts were taken to minimize the number of animals used and to reduce pain and suffering.

### Experimental Design

#### Animals

Male Long Evans rats weighing between 300 and 325 g were purchased from Harlan or Charles River. Rats were given one week to acclimate with daily handling prior to behavioral training or surgery and were then kept with regulated access to food to maintain 90% of their free-freeding body weight. They were given ∼18g of standard rat chow each day in the evenings following experiments. Rats were single-housed in their home cages in a 12h light/dark cycle colony room, with experiments occurring during the light cycle. A total of 11 rats had a microwire array implanted into medial frontal cortex. Some rats additionally had a drug cannula implanted into the opposite hemisphere using the same stereotaxic coordinates. Arrays were made by Tucker-Davis Technologies and consisted of 16 blunt-cut 50-µm tungsten wires, insulated with Formvar, separated by 250 µm within each row and 500 µm between rows. In vitro impedances for the microwires were ∼150 kΩ.

#### Surgeries

Animals had full access to food and water in the days prior to surgery. Stereotaxic surgery was performed using standard methods (e.g., Narayanan and Laubach, 2006). Briefly, animals were lightly anesthetized with isoflurane (2.5% for ∼2 min), and were then injected intraperitoneally with ketamine (100mg/kg) and either xylazine (10 mg/kg) or dexdomitor (0.25mg/kg) to maintain a surgical plane of anesthesia. Craniotomies were made above the implant location. Microwire arrays were placed into the medial frontal cortex (coordinates from bregma: AP: +3.2 mm; ML: + 1.0 mm; DV: -2.2 mm from the surface of the brain, at a 12° posterior angle). Four skull screws were placed along the edges of the skull and a ground wire was secured in the intracranial space above the posterior cerebral cortex. Electrode arrays were connected to a headstage cable and modified Plexon preamplifier during surgery and recordings were made to assess neural activity during array placement. Drug cannulas, 26-gauge PEEK (Plastics One), were implanted prior to the microwire arrays using similar procedures. Craniotomies were sealed using cyanocrylate (Slo-Zap) and an accelerator (Zip Kicker), and methyl methacrylate dental cement (AM Systems) was applied and affixed to the skull via the skull screws. Animals were given a reversal agent for either xylazine (yohimbine, 2mg/ml) or dexdomitor (Antisedan, s.c. 0.25 mg/ml) and Carprofen (5 mg/kg, s.c.) was administered for postoperative analgesia. Animals recovered from surgery in their home cages for at least one week with full food and water, and were weighed and handled daily.

#### Behavioral Apparatus

Rats were trained in operant chambers housed within a sound-attenuating external chamber (Med Associates, St. Albans, VT). Operant chambers contained a custom-made drinking spout that was connected to multiple fluid lines allowing for multiple fluids to be consumed at the same location. The spout was centered on one side of the operant chamber wall at a height of 5 to 6.5cm from the chamber floor. Tygon tubing connected to the back of the drinking spout would administer the fluid from a 60cc syringe hooked up to a PHM-100 pump (Med Associates). A “light-pipe” lickometer (Med Associates) detected licks via an LED photobeam, and each lick triggered the pump to deliver roughly 0.025 ml per 0.5 sec. Behavioral protocols were run though Med-PC version IV (Med Associates), and behavioral data was sent via TTL pulses from the Med-PC software to the Plexon recording system.

#### Continuous-Access Shifting Values Licking Task

The operant licking task used here is similar to that previously described in Parent et al. (2015a,b). Briefly, rats were placed in the operant chamber for 30 min, where they were solely required to lick at the drinking spout to obtain a liquid sucrose reward. Licks activated the syringe pumps to deliver liquid sucrose over 0.5 sec. Every 30 sec, the reward alternated between a high concentration (20% weight per volume) and low concentration (2-4% wt/vol) of sucrose. The animal’s licking behavior was constantly monitored.

#### Instrumental Shifting Values Licking Task

The operant licking task used above was modified slightly to allow for assessment of reinforced versus non-reinforced licks. A 2-sec inter-pump interval was included between each pump activation. In other words, the rat would lick to activate a liquid sucrose reward for 0.5 sec, and then once the pump stopped delivering fluid, no reward was delivered again for 2 sec. The next lick after the 2 sec interval would initiate the next pump activation. Licks during the 2 sec inter-pump interval were *instrumental*.

#### Multi-Electrode Recordings

Electrophysiological recordings were made using a Plexon Multichannel Acquisition Processor (MAP; Plexon; Dallas, TX). Local field potentials were sampled on all electrodes and recorded continuously throughout the behavioral testing sessions using the Plexon system via National Instruments A/D card (PCI-DIO-32HS). The sampling rate was 1 kHz. The head-stage filters (Plexon) were at 0.5 Hz and 5.9 kHz. Electrodes with unstable signals or prominent peaks at 60 Hz in plots of power spectral density were excluded from quantitative analysis.

#### Reversible Inactivation

Animals were tested with muscimol infusions in one hemisphere and recordings of neural activity in the opposite hemisphere. For control sessions, phosphate-buffered saline (PBS) was infused into MFC. The next day, muscimol (Sigma-Aldrich, St Louis, MO) was infused at 0.1 μg/μl. Infusions were performed by inserting a 33-gauge injector into the guide cannula, and 1.0 μl of fluid was delivered at a rate of 15 μl per h (0.25 μl per min) with a syringe infusion pump (KDS Scientific, Holliston, MA). The injector was connected to a 10 μl Hamilton syringe via 0.38 mm diameter polyethylene tubing. After infusion was finished, the injector was left in place for at least 4 min to allow for diffusion of the fluid. The injector was slowly removed and the headstage cable was subsequently plugged into the animal’s implant. Rats were tested in the instrumental Shifting Values Licking Task 1 hour after the PBS or muscimol infusions. Recordings were made the day following the infusion session without any manipulation to verify recovery from the inactivation session.

#### Histology

After all experiments were completed, rats were deeply anesthetized via an intraperitoneal injection of Euthasol (100mg/kg) and then transcardially perfused using 4% paraformaldehyde in phosphate-buffered saline. Brains were cryoprotected with a 20% sucrose and 10% glycerol mixture and then sectioned horizontally on a freezing microtome. The slices were mounted on gelatin-subbed slides and stained for Nissl substance with thionin.

### Statistical Analysis

#### Software and Testing

All data were analyzed using GNU Octave https://www.gnu.org/software/octave/), Python (Anaconda distribution: https://www.continuum.io/), and R https://www.r-project.org/). Analyses were run as Jupyter notebooks http://jupyter.org/). Statistical testing was performed using standard (base) packages for R. Standard non-parametric tests and paired t-tests were used throughout the study. Repeated-measures ANOVA (with the error term due to subject) were used to compare data across training sessions (Figure 7), reinforced versus non-reinforced licks (Figure 9), and PBS versus muscimol (Figure 10). Computer code used in this study is available upon request from the corresponding author.

**Figure 1:**
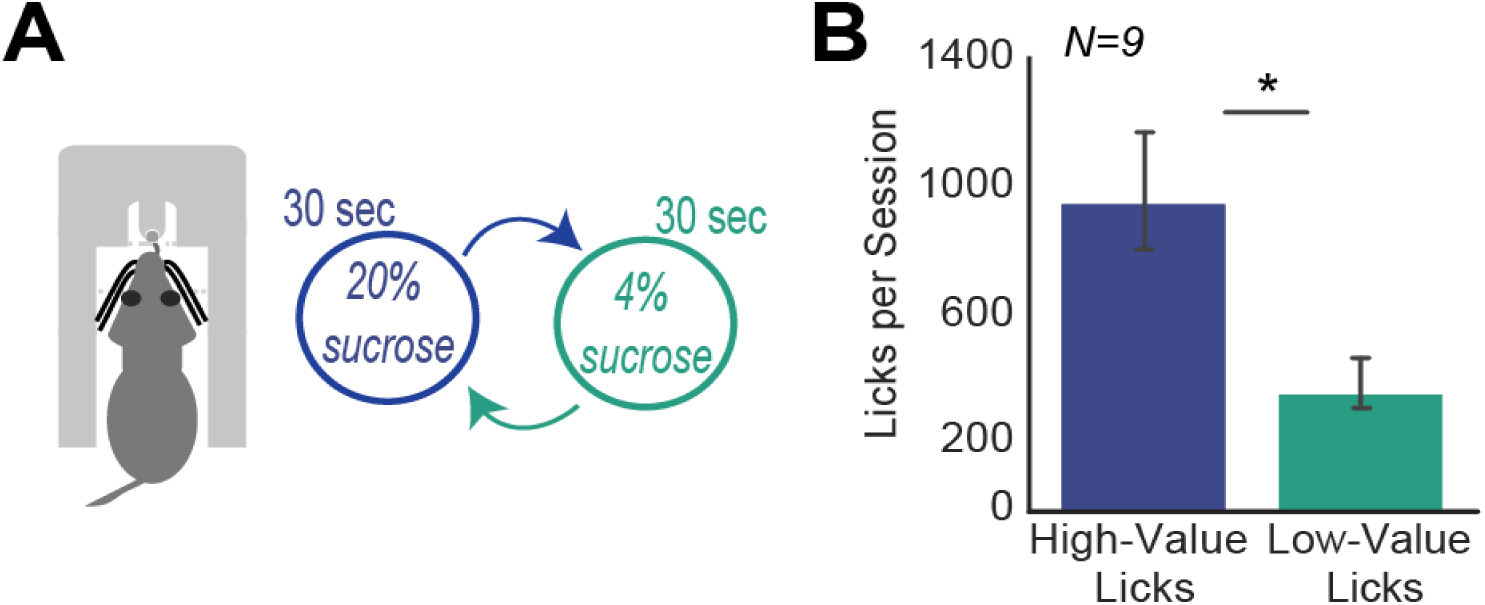
Behavioral task. A. Rats were tested in an incentive contrast procedure called the Shifting Values Licking Task (Parent et al., 2015a). They were required to lick on a spout to receive liquid sucrose rewards. Reward values shift between relatively high (20% wt/vol) and low (4% or 2% wt/vol) concentrations of sucrose every 30 sec. B. Experienced rats (fourth training session) licked more for the high-value sucrose than for the low-value sucrose (paired t-test; t(8) =4.29, p = 0.0026). \**p*<0.005.

**Figure 2:**
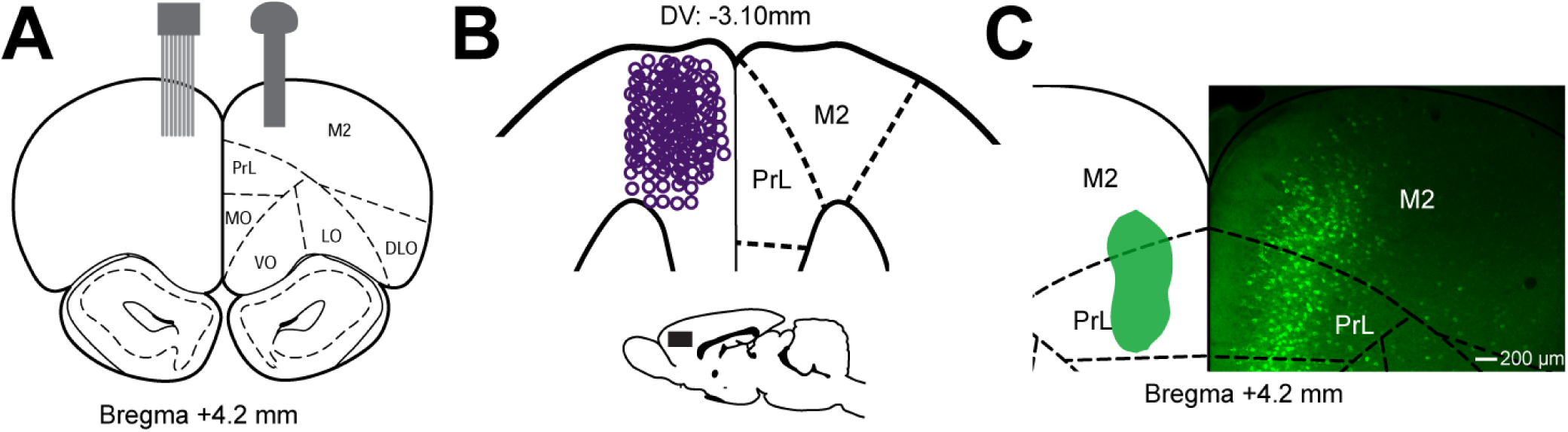
Neuronal recordings. A. All rats (N=11) were implanted with a 16-channel microwire array targeting the rostral medial frontal cortex [MFC] in one hemisphere. A subset of rats (N=4) had a drug cannula implanted in the same cortical area in the opposite hemisphere. B. Locations of recording sites are depicted on a horizontal section from the Paxinos and Watson (1997) atlas. All electrodes were placed within the prelimbic [PrL] and medial agranular [M2] regions. C. Validation of cross-hemispheric connections for this rostral MFC region. Cholera Toxin subunit B with the Alexa Fluor 488 reporter was injected in the rostral MFC of 5 rats. Injection site spread is schematically represented in green (left hemisphere). Neurons were labeled in the superficial layers in the opposite hemisphere (right).

**Figure 3:**
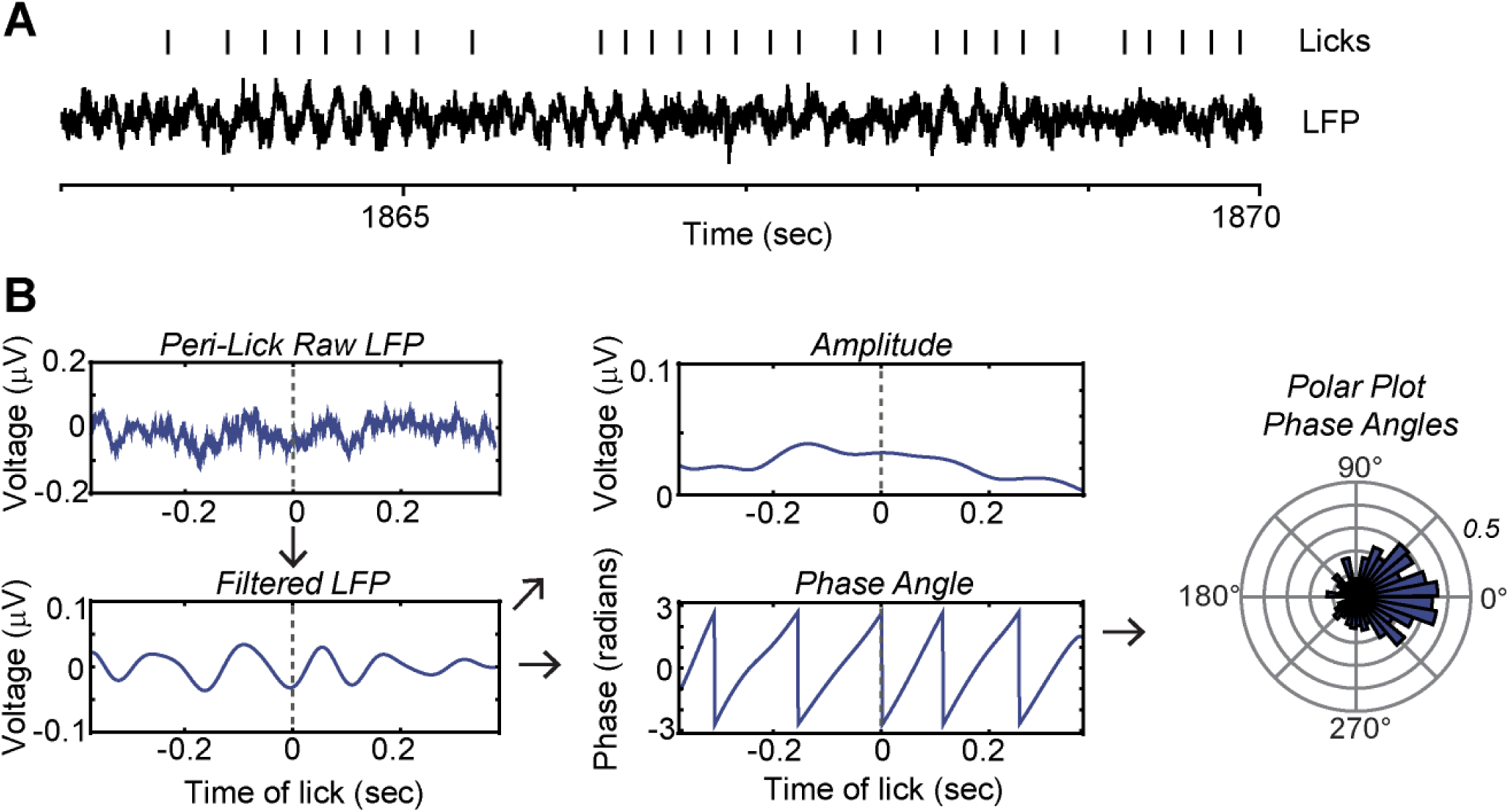
Neuronal activity in the MFC is entrained to the lick cycle. A. An example of a local field potential [LFP] recording shows clear fluctuations at the times of licks (tick marks above the LFP). B. Relationships between LFP signals and licking were assessed by bandpass filtering the LFPs near the licking frequency (defined by the inter-quartile range around the medial inter-lick interval) and applying the Hilbert transform to measure the amplitude and phase of licking-related neuronal activity. Instantaneous phase was plotted using polar coordinates and analyzed with standard methods for circular statistics (Agostinelli and Lund, 2013). See methods for details.

**Figure 4:**
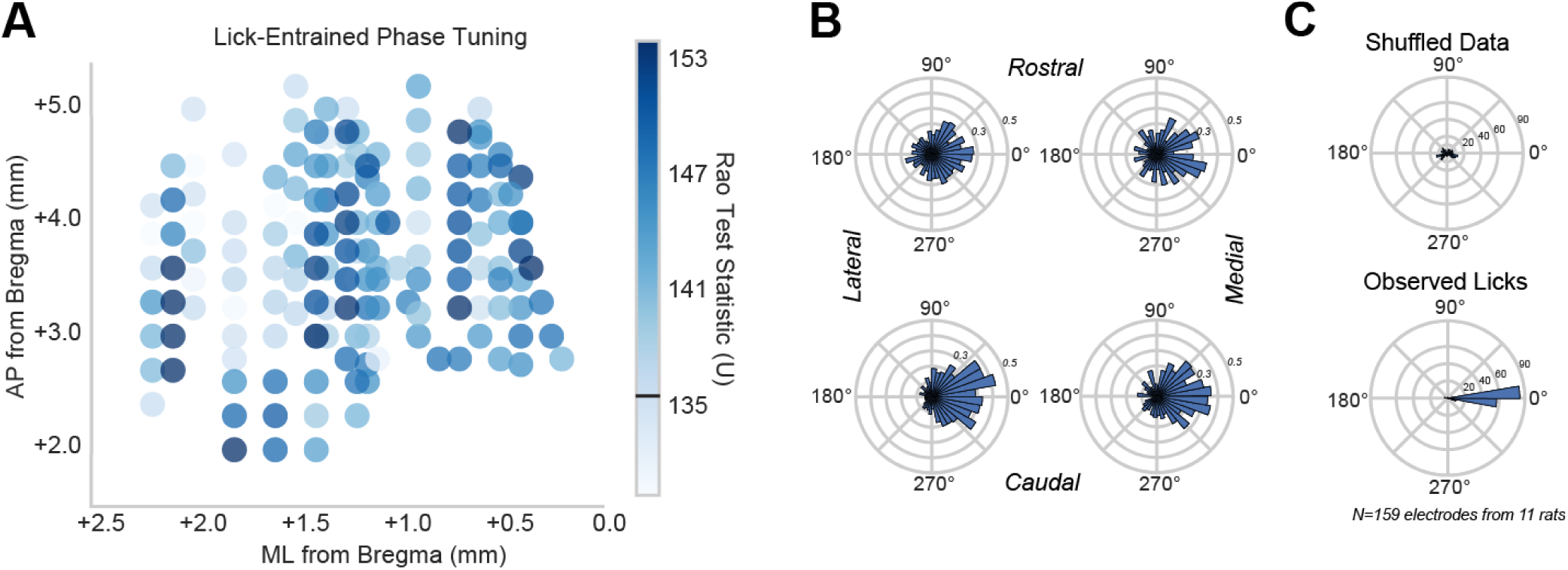
Spatial distribution of entrainment to the lick cycle. A. Spatial plot of phase tuning using the test statistic from Rao’s spacing test of uniformity showed no obvious topography of lick-entrainment in the MFC. Individual electrode locations were plotted according to their location in reference to Bregma (N=159 electrodes). Recording sites were depicted as circles colored by the strength of their Rao test statistic [U]. The colorbar shows values of U from the 5th to 95th percentile range over all recording sites. Values above the black bar (near 135) were not uniform (p<0.05). B. Polar plots represent phase tuning examples from four spatial extremes of the graph in (A). The most rostral/lateral (top left; U=134.48, p>0.05), rostral/medial (top right; U=152.30, p<0.001), caudal/medial (bottom right; U=153.51, p<0.001), and caudal/lateral (bottom left; U=147.44, p<0.001) electrodes were chosen. There was no drastic difference among the four locations with regard to phase tuning. C. Group summaries of the mean phase angle at the time of licking from all 11 rats reveal significant phase tuning toward 0 degrees (i.e., peak or trough of the rhythm). These results were compared with phase angles measured from surrogate data (shuffled inter-lick intervals), which did not show evidence for significant phase entrainment.

**Figure 5:**
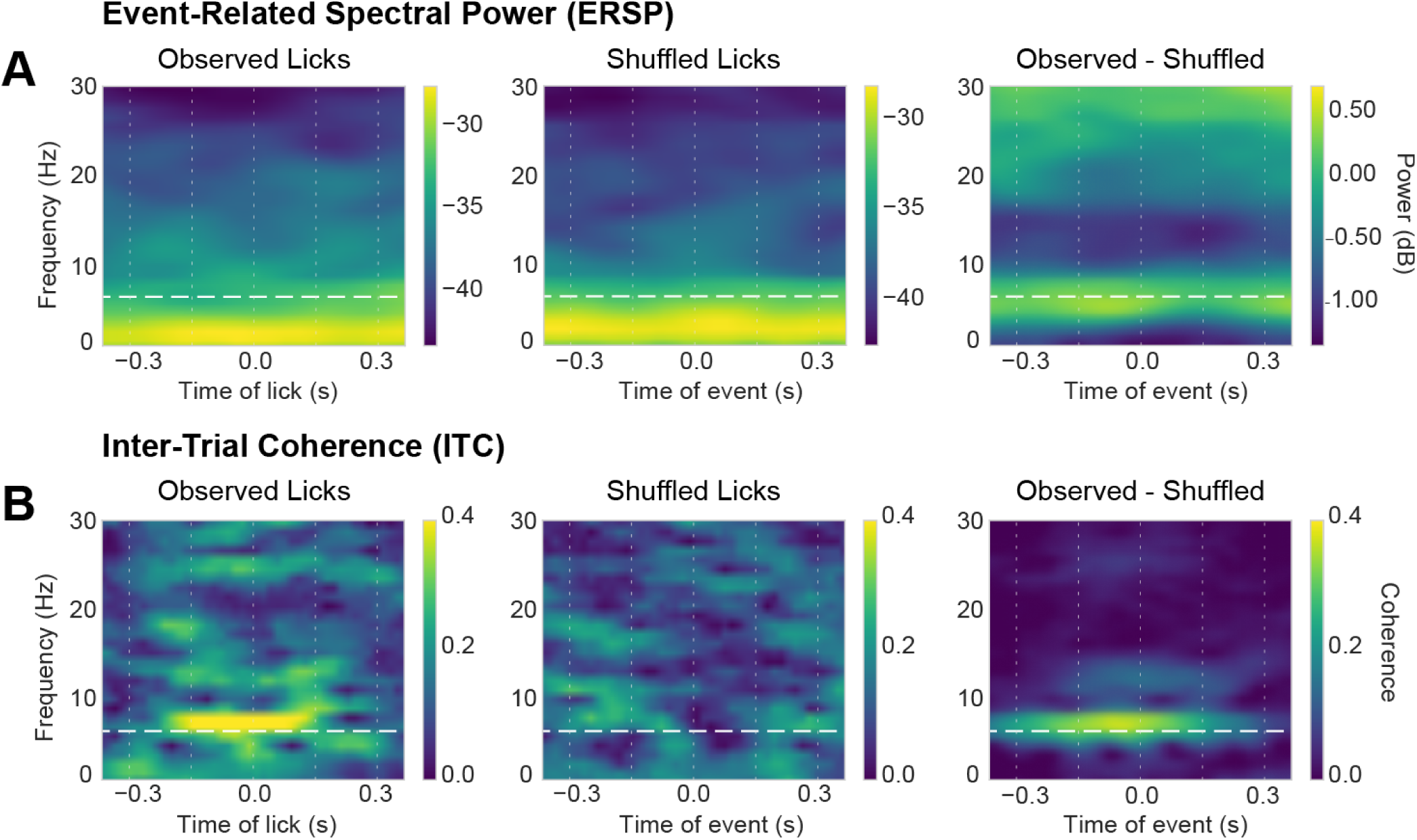
Time-frequency analysis of lick-entrained LFP data. Event-Related Spectral Power [ERSP] (top) and Inter-Trial Coherence [ITC] are shown for a typical LFP recording aligned to the time of licking in the behavioral task. The white horizontal dashed line depicts the median licking frequency. The white vertical dashed lines depict the median inter-lick intervals. ERSP and ITC measures were computed using observed licks (left) and surrogate data (middle), created by shuffling inter-lick intervals. A. Persistent elevated ERSP was notable at very low frequencies (∼2 Hz, or delta) for both the observed (upper left) and shuffled (upper middle) events, i.e., was not entrained to the lick cycle. Subtraction of the shuffled ERSP matrix from the observed ERSP matrix revealed elevated power at the licking frequency (horizontal dash line). B. ITC was apparent near the licking frequency over a period of two lick cycles for the observed licks (lower left), but not the shuffled licks ((lower middle). Subtraction of the shuffled ITC matrix from the observed ITC matrix revealed elevated power at the licking frequency (horizontal dash line).

**Figure 6:**
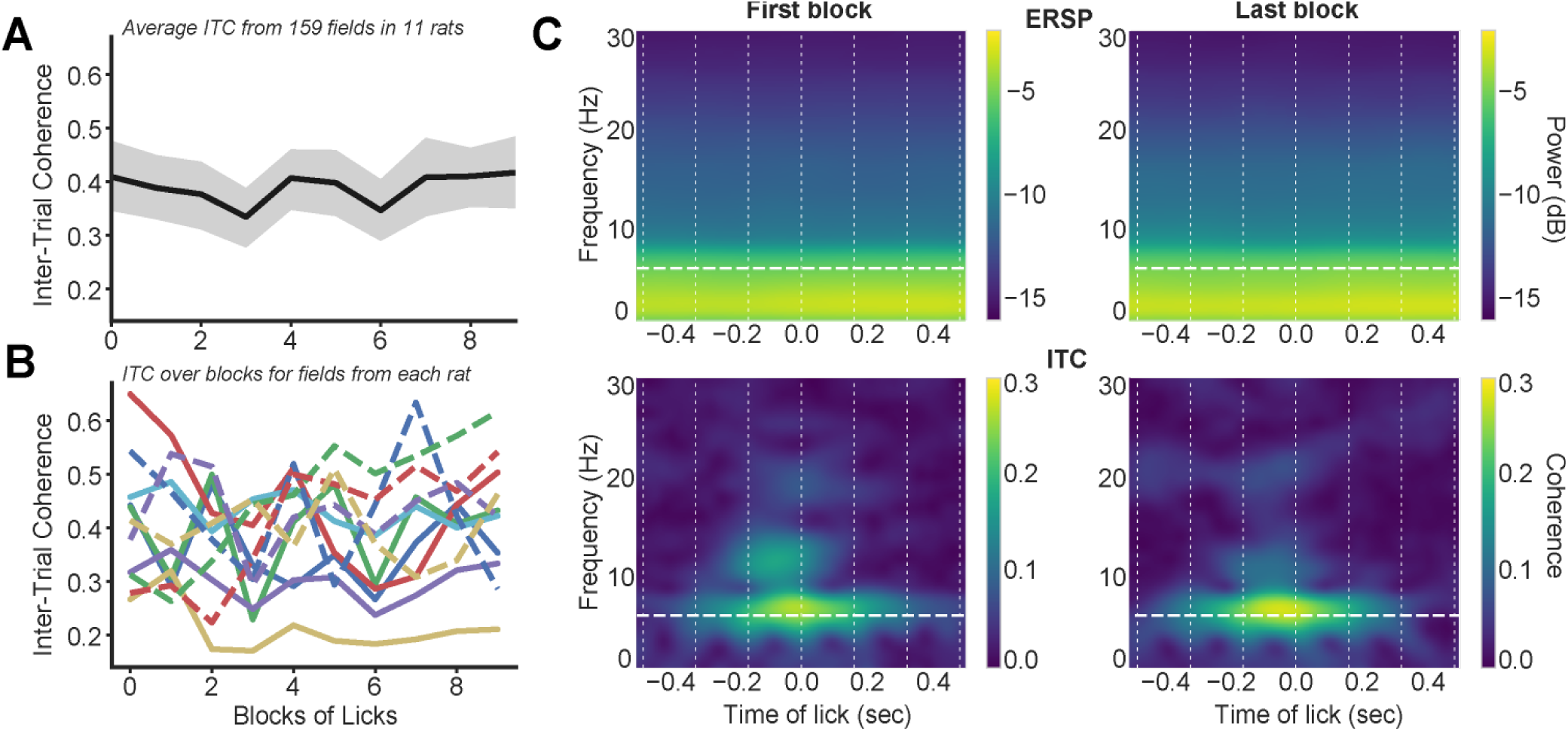
Entrainment was stable over the 30-min test sessions. Sessions were split into 10 blocks with equal numbers of licks and peak event-related spectral power [ERSP] and inter-trial coherence [ITC] were measured in the theta frequency range (6-12 Hz) over the inter-lick interval before and after each lick. A. Group average of peak ITC showed no evidence for a change in this measure over the data sets. Similar results were obtained for ERSP (not shown). B. Traces for peak ITC from each of the 11 rats. C. Grand average of ERSP and ITC for all LFPs in the first and last block. Together, these results suggest that entrainment of MFC LFPs to the lick cycle was not sensitive to cross-session factors such as satiety.

**Figure 7:**
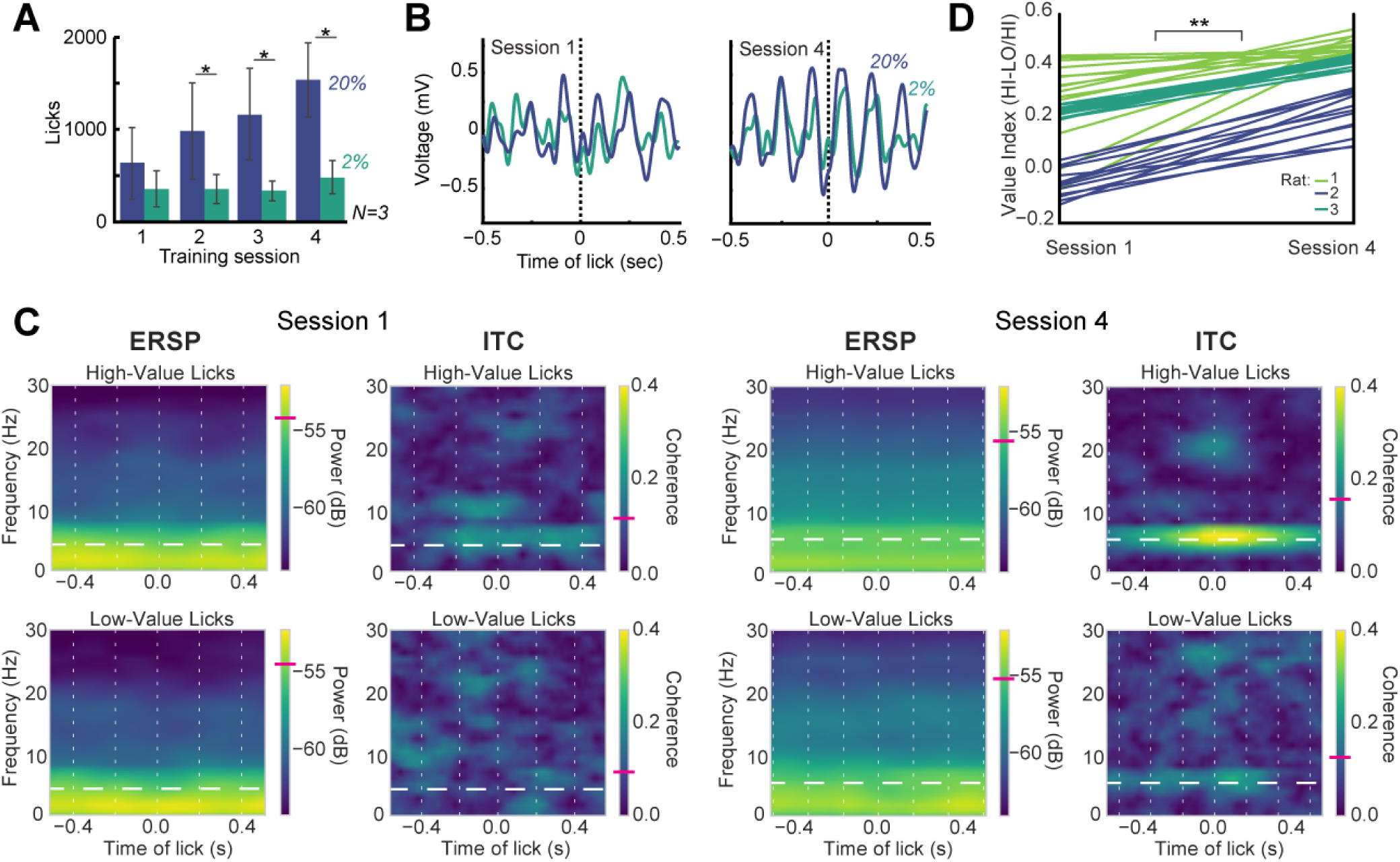
MFC theta entrainment to licking develops with experience. A. Recordings were made in a subset of three rats as they learned the behavioral task. The rats showed increased licking for the high-value sucrose compared to the low-value sucrose after the first training session and the relative difference in licking increased over the first four training sessions. B. Neuronal entrainment to the lick cycle developed with experience in the task. For example, event-related potentials increased in size and apparent rhythmicity between the first and fourth training session (blue = higher-value 20% sucrose; green = lower value 2% sucrose). C. Increased entrainment to the lick cycle was also apparent in Inter-Trial Coherence [ITC], which was not apparent in session 1 and specific to licks that delivered high-value sucrose in session 4. (White vertical lines = average inter-lick intervals across the session. White horizontal dashed line = average licking frequency across the session. Magenta ticks in the colorbars denote average ERSP or ITC at the median licking frequency.) D. To capture differences in ITC values for the two types of licks across all recordings, we used a value index, defined as ((ITC-HI – ITC-LO)/ITC-HI). The index was based on the peak ITC values in a temporal window ranging from one inter-lick interval before lick onset up to 50 ms after the lick and for all frequencies between 4 and 12 Hz (“theta”). As shown in the parallel line plot, in which each line denotes a LFP recording from a distinct electrode, this index was larger in session 4 compared to session 1 (paired t-test: t(39)=-12.085, p<10^-6^). *p<0.05; \*\**p*<10^-6^.

**Figure 8:**
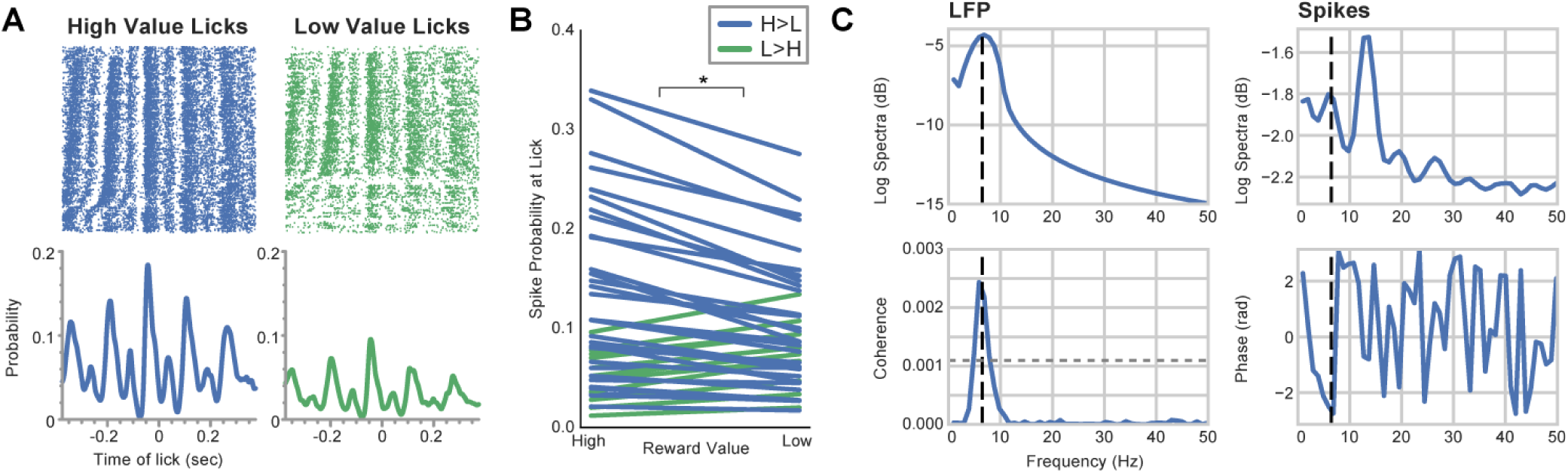
Coherence between spikes and licks reflects reward information. A. Multi-unit spike activity [MUA] was entrained to the lick cycle (high-value licks = blue; low-value licks = green). Rasters were sorted by the latency to the last lick before the lick at time 0, with the shortest preceding intervals at the top of the raster. The high-value licks were sub-sampled for this plot so that neural activity could be compared for the same number of total licks (at time 0). Peri-event histograms (bin: 1 ms, 10-point Gaussian smoothing), below the raster plots, denote the probability of spiking around the times of the licks. B. Group summary for spike probability at times of higher and lower value licks. Blue lines indicate higher spike probability for the higher value sucrose. Green lines indicate higher spike probability for the lower value sucrose. Spike probability was higher at the times of the higher value licks compared to times of the lower value licks (paired t-test: t(43)=3.78, p<0.001). 33 of the 44 MUA recordings showed higher spike probabilities for the higher value licks. C. Spike-field coherence found that all 44 MUA recordings were entrained to the LFP fluctuations that encoded reward information. Power spectra are shown in the upper row for example LFP and MUA recordings. Peak power was near the licking frequency (black dashed line) for the LFP. The main peak for the spike train was in the low beta range (12-15 Hz) and a second peak was at the licking frequency. Coherence between these signals (lower left plot) was found at the licking frequency (5.96 Hz), at a level approximately twice the 95% confidence interval. The phase between the spikes and fields at the licking frequency (lower right plot) was near–P, suggesting that the spikes and fields had an antiphase relationship. \**p*<0.001.

**Figure 9:**
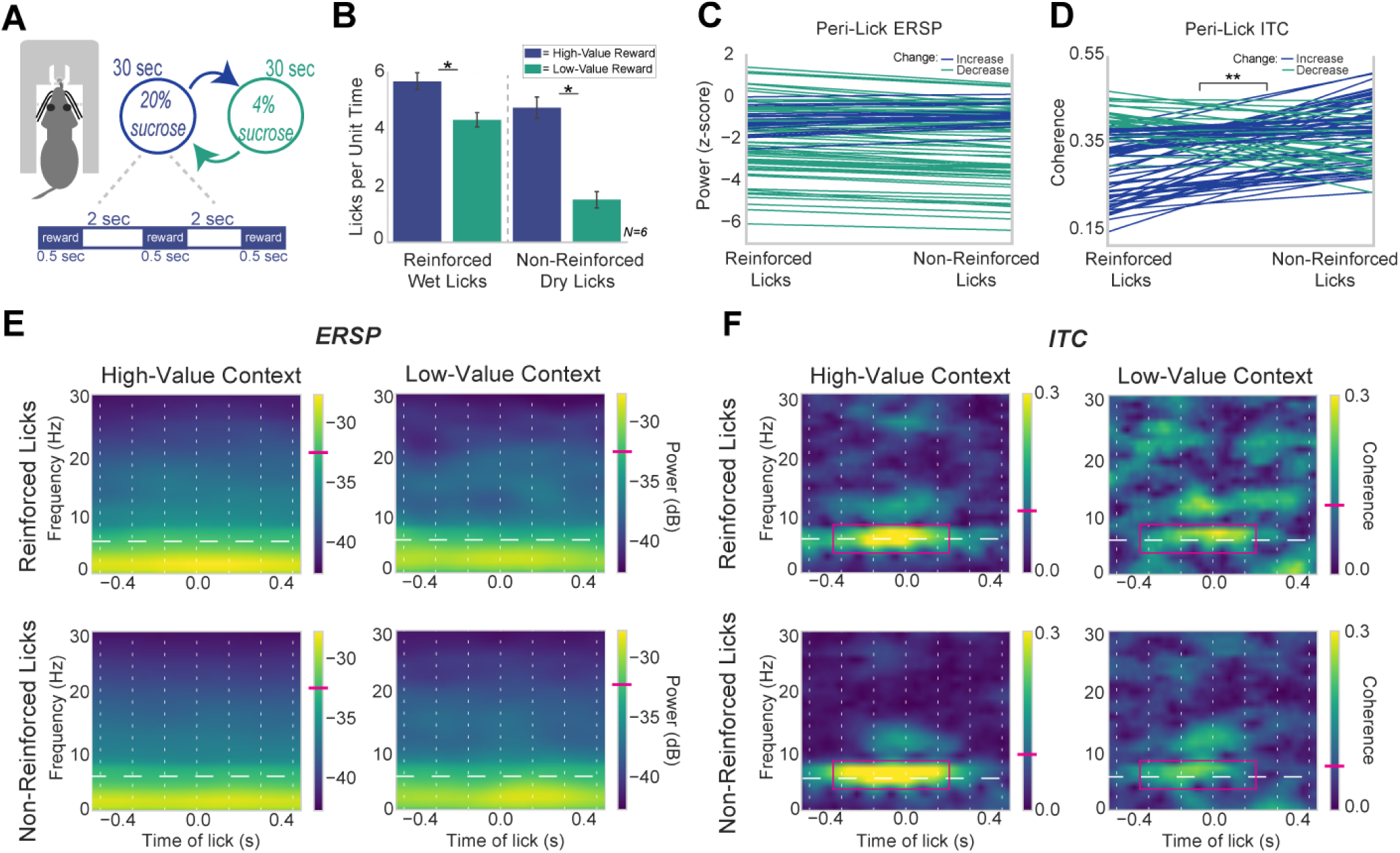
Reward context, not reinforcement per se, drives reward signaling. A. The Shifting Values Licking Task was modified to include a 2 sec period between reward deliveries. This period allowed for non-reinforced licks (dry licks at the spout) to be recorded within the 30 sec states of high or low-value sucrose availability. B. Group summary (N=6) of licks per unit time (total licks emitted in each context divided by time spend in each context). This measure revealed that rats licked less in the non-reinforced lower-value blocks compared to the other blocks. C. Peak ERSP values for reinforced versus non-reinforced licks during the high-value blocks. Lines are colored by their direction (increase or decrease in power). There was no difference in power for reinforced versus non-reinforced licks (F_(1,359)_=2.52, p=0.11). D. Peak ITC values for reinforced versus non-reinforced licks during the high-value blocks. The majority of LFPs showed increased phase-locking to non-reinforced licks (blue lines), while electrodes from two rats show a slight decrease in phase-locking for non-reinforced licks (green lines). Overall group summaries show an increase in phase-locking for the non-reinforced licks (F_(1,359)_ = 31.94, p<10^-6^). E,F. Example of time-frequency analysis of a LFP from a rat that showed decreased ERSP and ITC (magenta box) when the rat licked in the lower-value context. ITC was higher near the licking frequency when the higher value reward was available, regardless if the licks were reinforced or not. Horizontal white lines indicate the within-session licking frequencies. Vertical white lines indicate the inter-lick intervals for each session. Magenta ticks in the colorbars denote average ERSP or ITC at the median licking frequency. \**p*<0.05; \*\**p*<10^-6^.

**Figure 10:**
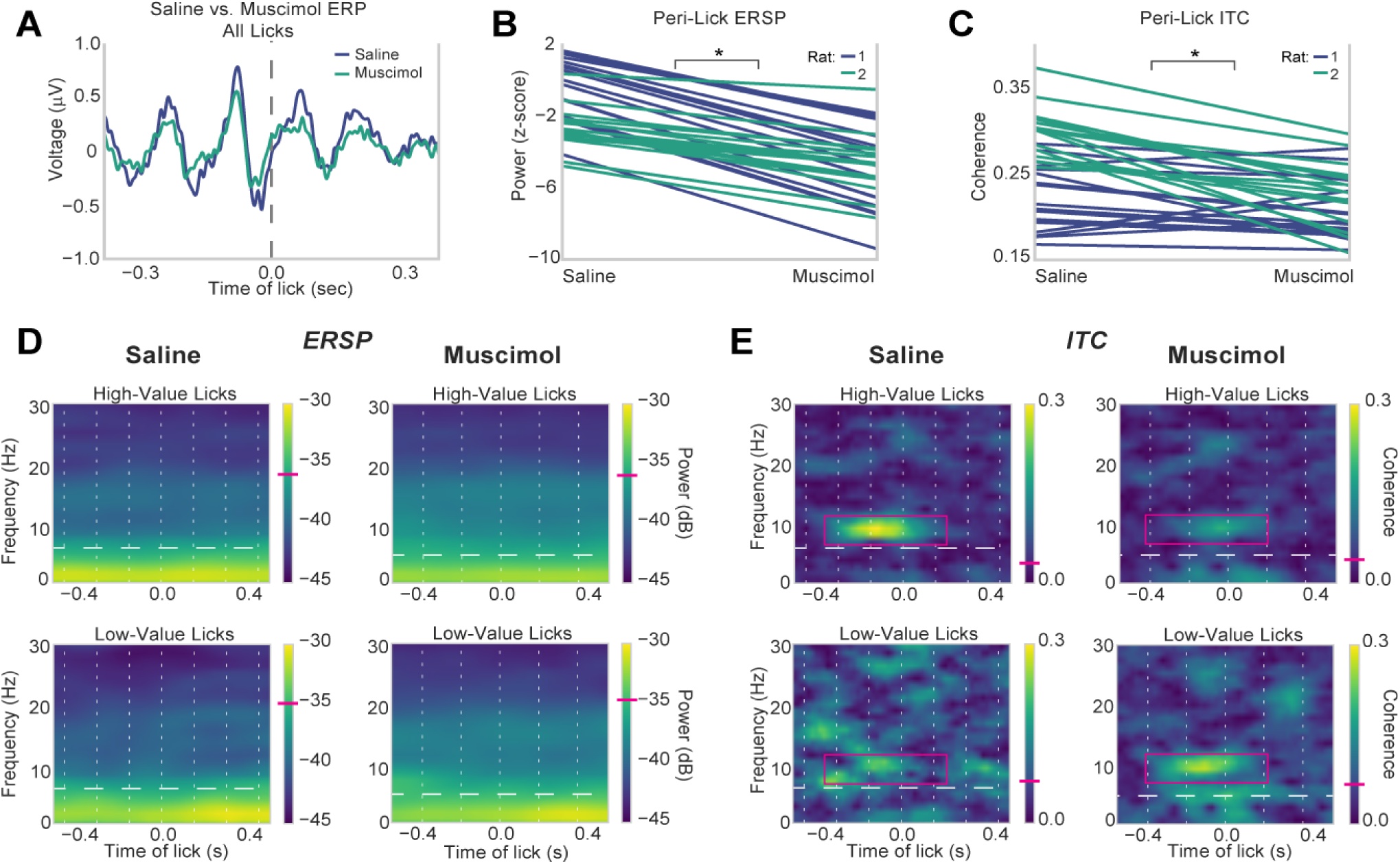
Reward signaling depends on neuronal activity in the MFC. Rats were tested with an electrode array in one hemisphere and an infusion cannula in the other, which was used to infuse either PBS or muscimol. A. Event-Related Potentials [ERP] from the saline (blue line) and muscimol (yellow line) sessions showed a similar overall time course around the licks. B. All electrodes showed a decrease in peak event-related spectral power [ERSP] at the licking frequency for higher value licks during the muscimol sessions compared to the PBS sessions (F_(1,123)_ = 96.09, p<10^-5^). C. Likewise, there was a reduction in peak inter-trial coherence [ITC] at the licking frequency for 28 of 32 electrodes (F_(1,123)_ = 18.17, p=3x10^-5^). D,E. Example of time-frequency analysis. Effects were specific to licks for the high-value reward. Horizontal white lines indicate the within-session licking frequencies. Vertical white lines indicate the inter-lick intervals for each session. Magenta ticks in the colorbars denote average ERSP or ITC at the median licking frequency. \**p*<10^-5^.

#### Data Analysis: Behavioral Measures of Licking

The average licking frequency for each rat was calculated by obtaining the inverse of the median inter-lick interval (ILI) across the behavioral session. Variability in this measure was estimated using the inter-quartile range.

To normalize the number of licks emitted in different reinforced or non-reinforced contexts that had different time lengths, licks per unit time (as represented in Figure 9B) were calculated by dividing the number of licks in each context by the actual amount of time spent in each context across the session. This was done by finding the sum of times between each context, and then subtracting from that time the amount of time of fluid delivery over the entire session.

Duration of licking bouts were detected, as in Parent et al., 2015a,b). Bouts of licks were defined as having at least three licks within 300 ms and with an inter-bout interval of 0.5 s or longer.

#### Data Analysis: Local Field Potentials

Electrophysiological data were first briefly assessed in NeuroExplorer http://www.neuroexplorer.com/). Subsequent processing was done using signal processing routines in GNU Octave. Analysis of LFP data was carried out using the EEGlab toolbox http://sccn.ucsd.edu/EEGlab/) (Event-Related Spectral Power and Inter-Trial Coherence) and Neurospec 2.0 http://www.neurospec.org/) (spike-lick and spike-field coherence). Circular statistics were calculated using the *circular* library for R (Agostinelli and Lund, 2013). Graphical plots of data were made using the *matplotlib* and *seaborn* library for Python. Analyses were typically conducted in Jupyter notebooks, and interactions between Python, R, and Octave were implemented using the *rpy2* and *oct2py* libraries for Python.

To measure the amplitude and phase of LFP in the frequency range of licking, LFPs were bandpass-filtered using EEGlab’s *eegfilt* function, with a fir1 filter (Widmann & Schröger, 2012), centered at the rat’s licking frequency (licking frequency + inter-quartile range; typically around 4 to 9 Hz), and were subsequently z-scored. Analyses were performed with a pre/post window of 2 sec, to capture effects at low frequencies, and the Hilbert transform was used to obtain the amplitude and phase of the LFP.

To measure the consistency of LFP phase angles in relation to licking, 500 licks were randomly chosen from one session from each rat along with 500 random time points that were chosen based on shuffling the inter-lick intervals from all licks in the rat’s session. After creating peri-event matrices from filtered and z-scored LFP data, the Hilbert transform was applied to obtain the phase angle and amplitude for each electrode, and analyzed with routines from the *circular* library for R. *rho.circular was used* to obtain mean resultant vector length. *mean.circular* was used to obtain average phase. The *rao.spacing.test* function was used to obtain Rao’s statistic and corresponding p-value, which indicates if phase angles were uniformly distributed.

For time-frequency analysis (ERSP and ITC), LFPs were preprocessed using EEGlab’s *eegfilt* function with a fir1 filter and bandpass filtered from 0 to 100 Hz. For group summaries, ERSP and ITC matrices were z-scored for that given rat after bandpass filtering the data. Peri-lick matrices were then formed by using a pre/post window of 2 sec on each side, and the *newtimef* function from the EEGlab toolbox was used to generate the time-frequency matrices up to 30 Hz. Plots of ERSP and ITC matrices were made using a narrow time window (∼0.5 sec) to focus on effects over several inter-lick intervals. Group summaries for ERSP and ITC were performed by obtaining the peak ITC value within a time window of ±2 interlick intervals (typically ∼±375 ms) around licking, and obtaining the peak ERSP value within that same window. Each electrode’s peak ERSP and ITC value for each type of lick (high-value or low-value lick) were used in the ANOVAs for group summaries. Finally, a “value index” was calculated to assess differences in ERSP and ITC measures associated with consumption of the higher and lower value rewards, i.e., (ITC_Hi_ – ITC_Lo_)/ITC_Hi_.

Shuffling methods were used to compare ERSP and ITC values for observed and shuffled licks (obtained by calculating inter-lick intervals, shuffling their trial order, and adding the intervals to the first lick in each behavioral session). This gave a set of surrogate licks with random timing unrelated to the animal’s behavior. Subsets of 50 licks and shuffled events were randomly chosen from each behavioral session and ERSP and ITC statistics were calculated for the subsets of observed and shuffled data. Shuffling was used to assess synchronous (theta) and asynchronous (delta) frequency ranges in the ERSP and ITC analyses (see Figure 5). However, statistical comparisons of ERSP and ITC values were made using raw spectral values.

Given the imbalance in the number of higher and lower value licks across the periods of learning and in the later testing sessions, all results were verified with subsampled sets of licks that were matched to the less represented condition (the square root of the less represented type) as well as using just 50 licks per type. Effects were consistent across these measures. Effects were also validated using the first 5 and 10 minutes of each testing condition, which were equivalent to effects across the entire session. To further assess the stability of the ERSP and ITC measures over the test sessions, the data sets were broken into 10 blocks with equal numbers of licks or equal amounts of time into the session. Peak ERSP and ITC values were calculated for frequencies between 4 and 12 Hz and within two inter-lick intervals around each lick. Summaries for each rat used grand average LFPs (z-scored). Plots of peak ERSP and ITC over the two types of blocks (licks, time) revealed no consistent cross-session effects, e.g., due to satiety (de Araujo et al., 2006; Bouret and Richmond, 2010), for both measures of neuronal activity (see Figure 6).

#### Data Analysis: Spike Activity

Exploratory analysis of on-line identified single units found that spike probabilities at the times of the licks were below 0.1 for all single units recorded in the task. Therefore, we used multi-unit activity [MUA] to relate spike activity to the animals’ lick cycles and related LFP signals. MUA was identified using the Plexon Offline Sorter v. 4.3 (Plexon, Dallas, TX). All recorded spike waveforms were thresholded (±2.7 times the standard deviation for the collection of waveforms) and “automatic artifact invalidation” was applied. Then, using routines in NeuroExplorer v. 5 (Nex Technologies, Madison, AL), we measured spike probabilities for all recorded MUAs around the higher and lower values licks, using 0.001 sec bins. Spike probabilities were compared for the two lick values using a paired t-test (in R). To measure Spike-Field Coherence [SFC], we also used routines (sp2a_m) from Neurospec 2.0 (*http://www.neurospec.org/*), and analyzed bandpass filtered LFP (licking frequency + inter-quartile range) and the following parameters: Segment power = 10 (1024 points, frequency resolution: 0.977 Hz), Hanning filtering with 50% tapering, and line noise removal for the LFPs at 60 Hz. To measure Spike-Lick Coherence [SLC], we used routines (sp2_m1.m) from Neurospec 2.0. The following parameters were used: Segment power = 12 (4096 points, frequency resolution: 0.244 Hz) and Hanning filtering with 50% tapering.

### Validation of cross-hemispheric connectivity using retrograde tracers

The methods for stereotaxic surgery that are described above were used to make injections of Cholera Toxin subunit B in 5 rats to validate cross-hemispheric connections within the rostral MFC, which have not been extensively studied in previous anatomical studies on the most rostral part of this cortical region (cf. Gabbott et al., 2003; Hoover and Vertes, 2007). The CTB had an Alexa Fluor 488 reporter from Molecular Probes and was injected at a 1% concentration and 400 nl volume. A 10 µl glass Hamilton syringe and Narishige motorized microinjector (IMS-10) was used. 10 min was allowed for diffusion after each injection. Brains were extracted using the methods described above for euthanasia and perfusion. Cortical slices were cut in the frontal plane using a freezing microtome and were imaged on fluorescent microscope (BX-51-F, Tritech Research, Los Angeles, CA) using an R1 camera and Ocular software from Qimaging (Surrey, BC).

## Results

### Multi-electrode recordings in the Shifting Values Licking Task

To investigate the role of the frontal cortex in reward-related consummatory behaviors, we assessed licking behavior in rats while performing simultaneous recordings in the rostral MFC. We trained rats in the Shifting Values Licking Task (Parent et al., 2015a), in which they licked at a drinking spout to receive 0.025 ml of a liquid sucrose reward (Figure 1A). The reward value of the fluid switched between higher (20% sucrose wt/vol) and lower (2 or 4%) levels every 30 sec. Experienced rats (>3 training sessions) licked more for the high-value reward compared to the low-value reward (Figure 1B; paired t-test between high-value and low-value licks: t_(8)_ =4.29, p = 0.0026).

Eleven rats were implanted with multi-electrode arrays (Figure 2A). The placement of electrodes is shown in Figure 2B. All electrodes were placed in the medial agranular and prelimbic areas. In four of the rats, a drug cannula was also implanted in the opposite hemisphere using the same stereotaxic coordinates. These animals were used to examine effects of reversible inactivation of the MFC without shutting down local neuronal activity. To confirm that cross-hemispheric connections exist within the rostral MFC region, 5 rats were injected with Cholera Toxin subunit B in the region where the neuronal recordings and reversible inactivations were made (Figure 2C).

### Quantification of lick-entrained rhythmic activity

We recorded 161 local field potentials [LFPs] from the MFC in 11 rats as they ingested liquid sucrose in the Shifting Values Licking Task. An example of licking-related fluctuations in the LFPs is shown in Figure 3A. To measure entrainment between LFPs and the animal’s licking, we bandpass filtered the LFPs around the licking frequency and used the Hilbert transform to extract the amplitude and phase of the peri-lick rhythm (left plots in Figure 3B). The phase of LFPs was plotted using polar histograms (right plot in Figure 3B).

To quantify relationships between licking and LFP phase, we used circular statistics to measure the consistency of the phase angles at the time of licks. We used Rao’s spacing test for uniformity, which assesses the directional spread of circular data. Two LFPs did not have major power in the 6-12 Hz range (see next page) and were excluded from this analysis. We plotted each electrode’s location in MFC and shaded the locations by the intensity of the Rao statistic (Figure 4A). Phase entrainment to the lick cycle was concentrated in three longitudinal zones within the prelimbic cortex (0 to 1 mm lateral the midline), the medial agranular cortex (“M2”) (1 to 1.5 mm lateral to the midline), and a the border of the medial and lateral agranular cortex (2 to 2.5 mm lateral to the midline). Examples of entrainment at four electrodes (from four different rats) located in each extreme of MFC space (rostral/lateral, rostral/medial, caudal/lateral, and caudal/medial) are shown in Figure 4B.

Remarkably, the LFPs as a population had a mean phase angle near 0 degrees, i.e., at the peak or trough of the neural oscillation (Figure 4C). This distribution was entirely distinct compared to population summaries based on surrogate data (i.e., “licks” derived from shuffled inter-lick intervals). Therefore, the MFC region as a whole showed a relatively similar phase that is suggestive of phase entrainment to the lick cycle.

To examine effects of licking on MFC across spectral frequencies, we used standard time-frequency analysis measures (EEGlab toolbox: Delorme and Makeig, 2004). Lick-related changes in spectral power across frequencies between 0 and 100 Hz were measured using Event-Related Spectral Power [ERSP] (Figure 5A). Phase consistency at the times of the licks was measured using Inter-Trial Coherence [ITC] (Figure 5B). For each LFP recording, we measured the frequency with the highest level of ERSP and ITC. Most LFPs (127 of 161) showed peak ERSP in the delta range (1-4 Hz). Most LFPs (104 of 161) showed peak ITC near the licking frequency, between 5.8 and 7 Hz. Some LFPs showed peak ITC near 8 Hz (N=27) or between 8 and 12 Hz (N=28). The LFPs with higher frequencies for peak ITC were recorded on common arrays, and our finding of variability among peak ITC frequencies might reflect differences in cortical layer or field, but we were unable to address these issues completely in the present study. The vast majority of LFPs (159 of 161) showed peak ITC at the time of licking between 5.8 and 12 Hz, and so we describe this frequency range as 6-12 Hz theta throughout the manuscript.

To determine if the spectral ERSP and ITC measures were synchronous with the lick cycle, or simply elevated during periods of licking but not time-locked to the actual licks (asynchronous), we created surrogate data by shuffling inter-lick intervals. This analysis provided evidence for both measures, ERSP and ITC, being elevated at the licking frequency in shuffle-corrected plots of ERSP and ITC (right panels in Figure 5A,B). This finding, together with the analysis using circular statistics described above, is strong evidence for an entrainment of 6-12 Hz theta activity in the MFC being entrained to the lick cycle.

As a previous study reported that consummatory-related activity in the MFC is sensitive to satiety (Bouret and Richmond, 2010; see also de Araujo et al., 2006 for related findings on the rodent orbitofrontal cortex), we examined if there were cross-session effects within our data sets that could reflect effects of satiey. We checked for stability in licking-related ERSP and ITC levels over the behavioral sessions by dividing licks into 10 blocks with either equal numbers of licks or equal amounts of time in the session. Peak levels of ERSP and ITC between 4 and 12 Hz and within one inter-lick interval around each lick were measured over the blocks. No obvious pattern indicating satiation was apparent in this analysis (ITC results are shown in Figure 6). This analysis suggests that entrainment of MFC activity to the lick cycle does not reflect satiety or other cross-session factors.

### Theta entrainment to licking develops with experience

In a subset of three rats, we recorded neuronal activity as the animals learned to perform the Shifting Values Licking Task (Figure 7). Over the first four days of training, the rats showed increased licking when the higher value reward was available relative to licking for the lower value option (Figure 7A). Repeated measures ANOVA found a main effect of reward value on licking (F_(1,14)_=32.20, p=5.7x10^-5^). Tukey’s *post hoc* test found evidence for a difference between the number of licks for the high-value versus low-value reward in session 4 (p=0.013), but not session 1 (p=0.935). Median inter-lick intervals (ILIs) were reduced from session 1 to session 4 (Wilcoxon rank-sum test from three rats individually: p<10^-6^), but this effect was relatively minor. The average median ILI in session 1 was 0.174+/-0.133 sec. The average median ILI in session 4 was 0.153+/-0.026 sec. This difference resulted in an increase of no more than 1.12 Hz in the average licking frequency between session 1 and session 4.

Entrainment to licking developed over the training sessions, with clear lick-related oscillatory patterns apparent in the LFPs by the fourth training session (Figure 7B). Event-related potentials were larger for licks that delivered the higher value sucrose reward (blue lines in Figure 7B). The spectral content of the signals was evaluated using the same ERSP and ITC analyses used in Figures 5 and 6. Notably, there was a distinction between high-value and low-value phase-locking to the onset of licking evident in LFP data from session 4, but this signal was not apparent during session 1 (Figure 7C). To capture effects of the different sucrose concentrations on the LFPs, we calculated a “value index” for each electrode (Figure 7D). This index was derived from difference between the peak ITC level for the high and low-value licks divided by the peak ITC level for the high-value licks. All electrodes from all rats showed an increase in this index, a result that indicates increased differences in phase-locking for two amounts of reward over the period of training (paired t-test: t(39)=-12.085, p<10^-6^).

To further measure changes in the signals associated with the two reward values over sessions, we performed a repeated-measures ANOVA with the peak ITC values as the dependent variable and the values of the licks and the training sessions as predictors. This analysis found a significant interaction between session and value (F_(1,155)_=22.43, p<10^-6^), and Tukey’s *post hoc* test found evidence for differences between session 1 versus session 4 high-value lick ITC values (p=0.0016). While these analyses found evidence for significant difference between ITC levels for the high and low-value licks in session 1 (p=0.0093), the difference between ITC levels was much greater in session 4 (p<10^-6^).

We also assessed changes in LFP power by performing the same type of repeated-measures ANOVA, using peak ERSP values as the dependent variable. There was a significant interaction between session and reward value (F_(1,155)_=9.991, p=0.0019). Tukey’s *post hoc* analyses showed a difference in power from session 1 to session 4 high-value licks (p=3.5x10^-4^), as well as power for session 4 high and low-value licks (p=4.3x10^-5^). There was no difference in power between session 1 high-value and low-value licks (p=0.99). These findings are evidence for learning-dependent changes in the theta entrainment by the event-related power (ERSP) and phase (ITC) to the lick cycle.

### MFC spike activity is entrained to the lick cycle

Simultaneous recordings of spike activity in the three rats tested during the final learning session showed evidence for spike entrainment to the lick cycle (Figure 8A). The probability of spiking at the times of licks was below 0.1 for all isolated single-units. Therefore, we re-isolated multi-unit activity [MUA] (N=44) for the analyses reported here and measured the probability of spiking at the times of the higher-value and lower-value licks. Spikes were more likely to be coincident with the higher-value licks (0.113±0.013, mean±sd) compared to the lower-value licks (0.092±0.009) for 33 of 44 spike recordings (Figure 8B; paired t-test: t(43)=-3.78, p=0.00047).

Spike-lick coherence was used to further analyze synchronization between spikes and the higher and lower value licks. Results were complicated (and thus not shown graphically), with spikes having major peaks at various frequencies in the beta and gamma ranges, and spike-lick coherence often being significant in those ranges. 33 of 44 units (75%) fired in phase with the higher value licks. 19 units (43%) fired in phase with licks that delivered the lower value fluid. Over all recordings, the level of spike-lick coherence was greater for the higher value licks compared to the lower value licks (paired t-test: t(43)=4.6, p<0.001).

A much simpler result was obtained by using spike-field coherence to examine synchronization between the spikes and fields (Figure 8C). All 44 MUA recordings exhibited significant levels of spike-field coherence at the licking frequency (lower left plot in Figure 8C). Interestingly, phase was uniformly near-P for these datasets (lower right plot in Figure 8C), indicating that spikes and fields were explicitly out of phase (antiphase). Together, these results suggest that the lick-entrained theta rhythmic activity as measured in the LFPs was also manifest in lick-entrained spike activity within the rostral MFC.

### Reward context, not reinforcement, drives licking-related theta entrainment

The signals described above could reflect the expected reward magnitude (van Durren et al., 2008) and/or the taste or fluid properties of the ingested solutions (Jezzini et al., 2013). To examine these issues, we modified the Shifting Values Licking Task to include a 2-sec period of non-reinforced licking between periods of reward delivery (Figure 9A). This procedure resulted in rats continuously licking at the spout during the non-reinforced blocks of the task. Six rats were tested in the procedure with neuronal recordings, and the occurrence of licking in each context (Licks per unit time) was analyzed by calculating the number of licks in each context divided by the actual time spent in each context across the session (Figure 9B). All rats continued to lick more during these non-reinforced blocks when they could receive the high-value fluid compared to when they could receive the low-value fluid (t_(5)_ = 5.20, p=0.0003 for all high-value context licks against all low-value context licks; t_(5)_=5.63, p=0.0025 for non-reinforced high-value context licks versus non-reinforced low-value context licks; t_(5)_=3.31, p=0.0213 for reinforced high-value context licks versus reinforced low-value licks).

LFP signals synchronized to reinforced and non-reinforced licks were similar, with the main differences between high-value licks and low-value licks still evident, despite the rats not being rewarded during the non-reinforced blocks. Figures 9C and D show group summaries of the differences in peak ERSP and ITC values at the onset of the reinforced and non-reinforced high-value licks. (We chose to focus on the high-value licks for the analyses due to the increased number of high-value licks emitted during the task, though low-value licks also show the same effect.) There were no major differences in peak ERSP levels for reinforced and non-reinforced licks (F_(1,359)_=2.52, p=0.11), which is also evident in the spectral plots from an example LFP recording shown in Figure 9E.

However, the majority (60 of 91) of the electrodes (from all rats) showed increases in ITC phase-locking values for the non-reinforced high-value licks (Figure 9D). We performed a repeated-measures ANOVA with factors for lick type (reinforced or non-reinforced) and reward value (high or low) with peak ITC values as the dependent variable. This analysis found evidence for a significant interaction between lick type and reward value (F_(1,359)_ = 31.94, p < 10^-6^). The non-reinforced licks had slightly greater ITC values at the onset of licking (high-value reinforced licks = 0.48, SD = 0.069; high-value non-reinforced licks = 0.51, SD = 0.063), which was also confirmed using Tukey’s *post hoc* test (reinforced versus non-reinforced high-value licks, p=0.0002). Spectral plots, shown in Figure 9F, revealed modest increases in phase-locking for the non-reinforced high-value licks, and minimal differences in the phase-locking for the reinforced versus non-reinforced low-value context licks.

These findings cannot easily be explained by how often the rats licked in each task context. If engagement in licking (vigor or intensity) explained the pattern of neural results, then we would have expected to find elevated ITC levels when the rats received liquid sucrose as well as when they made non-reinforced licks in the high-value blocks. However, ITC levels were not elevated when rats licked for the lower concentration of sucrose. We examined two other behavioral measures of licking to determine if the differences in ITC levels could be accounted for by a behavioral measure from the task. First, we examined the licking, frequency (based on inter-lick intervals) but found no differences between median inter-lick intervals (Mann-Whitney test: U_5_=-0.3202, p=0.7487) and the inter-quartile ranges for the inter-lick intervals (U_5_=-1.4412, p=0.1495) in the higher and lower reward contexts, findings that discount the role of the frequency of licking in explaining the ITC results. By contrast, when we examined the persistence of licking, using bout analysis (as in Parent et al., 2015a,b), we found clear differences in the duration of licking bouts in the higher (5.8003±0.9278 sec) and lower (1.6969±0.3932 sec) reward contexts (U_5_=2.5621, p=0.0104). Bouts were on average ∼3.4 times longer in the higher value contexts. As such, the differences in ITC levels across the higher and lower reward contexts were associated with the persistence with the rats licked in the two behavioral contexts, but not the vigor or frequency at which they licked. These findings suggests that reward expectation, rather than the properties of the delivered fluids, drives reward signaling in the rostral MFC.

### Reward signaling depends on the medial frontal cortex

Having established the MFC as signaling reward information through lick-entrained neuronal rhythms, we carried out a reversible inactivation study to determine if the rhythmic activity is generated by neurons in the MFC. (Alternatively, the signals could be generated elsewhere and broadcast to the MFC.) We implanted four rats with a multi-electrode array in one hemisphere and a drug infusion cannula to allow for infusing muscimol in the MFC in the opposite hemisphere (Figure 2A). Cross-hemispheric inactivation was done to allow for recording distant effects of MFC perturbation, and not the effects of a local shutdown of neuronal activity. Two of these rats had precisely aligned electrode arrays and drug cannula (same cytoarchitectural area and layer). Two other rats were not precisely aligned (e.g., cannula in superficial layers and array in deep layers), and electrophysiological data from those animals were not considered further. In all four rats, we did not observe any major behavioral change in the number of licks emitted after the unilateral muscimol infusions. There was a marginal decrease in the overall inter-lick intervals under muscimol (Mann-Whitney U test, p<0.05). There was no difference in bout durations for the high or low value licks between the saline and muscimol sessions (p>0.05 for all comparisons, Mann-Whitney U test). (This is in contrast to our previous study with bilateral inactivations (Parent et al., 2015a), which clearly alter performance of the task, including the persistence of licking bouts.) The lack of behavioral effects of muscimol allowed us to assess potential electrophysiological changes without overt effects of the inactivations on the animals’ licking behavior.

In the two rats with aligned electrode arrays and drug cannulas, LFP activity during muscimol inactivations was dramatically altered. Muscimol infusions slightly decreased event-related potentials synchronized to licking (Figure 10A) and dramatically decreased event-related spectral power [ERSP] at the licking frequency (Figure 10B, all electrodes plotted from two rats). This was confirmed in the spectral plots, shown from one example electrode from one rat (Figure 10D). A repeated measures ANOVA on peak ERSP values around the onset of licking found evidence for decreased power at lick-onset between the saline and muscimol sessions (F_(1,123)_ = 96.09, p<10^-6^). Muscimol infusions also decreased the lick-entrained phase-locking in the theta frequency range. As seen in Figure 10C, 28 of 32 electrodes showed decreased phase-locking around the onset of licking. A repeated measures ANOVA found evidence for difference in ITC values for the saline and muscimol sessions (F_(1,123)_ = 18.17, p=3.9x10^-5^). Spectral plots from an example electrode (Figure 10E) show diminished phase coherence in the theta frequency range for the high-value licks. The decrease in phase-locking therefore disrupted the previously evident differential signaling of high and low value sucrose rewards. Together, this inactivation study established that the theta-entrained activity described throughout this study, and also reported for single-value water rewards in Horst and Laubach (2013), are generated by neurons in the rostral MFC.

## Discussion

### Dynamics of reward-related activity in the MFC

The main idea of this study was that rodents lick to consume rewarding fluids and reward information is therefore likely to be encoded in a dynamic manner, in phase with the rewarded actions. Initial evidence for this idea was reported by Horst and Laubach (2013), who found MFC neurons that were selectively activated when thirsty rats initiated bouts of licking to consume water. The present study was designed to further examine the dynamics of reward-related activity in the MFC as rats ingested varying levels of liquid sucrose during a MFC-dependent incentive-contrast licking task (Parent et al., 2015a,b). Multi-electrode recordings were made in the rostral MFC as rats consumed relatively higher and lower value rewards that were available in alternating periods of 30 sec (Figures 1 and 2). We found that the entrainment of MFC spikes and field potentials to the animals’ lick cycle varied with the concentration of liquid sucrose that was ingested (Figures 3-6 and 8). These signals were distributed broadly throughout the rostral part of the MFC (Figure 4A). Spectral methods showed that these effects were selective to the 6-12 Hz theta range, which also encompasses the animals’ licking frequency (right plots in Figure 7C). We further examined if theta-entrained activity to the lick cycle is stable across the testing sessions (Figure 6), develops with experience (Figure 7), does not depend on the presence of the rewarding fluids (Figure 9), and depends on processing by MFC neurons (Figure 10).

The differential expression of lick-entrained MFC theta might reflect the reward value of the ingested fluids and/or the vigor or persistence with which rats lick to consume the fluids. As the rostral MFC is known to have extensive interconnections with the “gustatory” insular cortex (Gabbott et al., 2003), MFC reward coding might reflect differences in the tastes or fluid properties of the sucrose solutions. Alternatively, the rostral MFC projects to a number of orolingual motor areas (Yoshida et al., 2009; Haque et al., 2010; Iida et al., 2010) and could mediate cortical control over the vigor (intensity per unit time) or persistence (bout structure) of licking. To examine these issues, we modified the task to have periods of non-reinforced licking between each reward delivery (Figure 9). If the presence of fluids drives MFC signaling, then differences in MFC activity should only occur when the animals were actively ingesting the fluids. Surprisingly, this experiment revealed that the theta-entrained activity persisted beyond the period of reward delivery (Figure 9), and was selectively elevated when the rats licked in the high-value blocks of the task without regard to reinforcement. These neuronal signals were associated with differences in the duration of licking bouts, with bouts being ∼3.4 times longer in the high-value blocks compared to the low-value blocks (see Results). The duration of licking bouts is classically interpreted as a measure of the palatability, or subjective value, of rewarding solutions (Davis and Smith, 1992). However, in our task design, the intervening periods of non-reinforced licking did not deliver liquid sucrose to the rats. Therefore, we suggest that the elevated MFC theta associated with more persistent licking in the higher-value blocks might have reflected the animal’s expectation of the higher-value reward. This interpretation should be verified in future studies, e.g. using shifts in sucrose concentration and fluid volume, which have opposing effects on the duration of licking bouts (Spector et al., 1998; Kaplan et al., 2001).

While the behavioral mechanisms mediating lick-entrained MFC theta will require new experiments to be resolved, our study was able to determine that the signals depend on MFC neurons. Reversible inactivation of the MFC (using muscimol) found evidence for MFC neurons being necessary for the generation of the lick-entrained signals (Figure 10). Perturbations of the MFC, made by infusing muscimol into the MFC in opposite hemisphere, dramatically attenuated lick-entrained MFC theta in the opposite hemisphere, directly implicating MFC neurons in the generation of the signals.

Together, our findings provide evidence for the MFC processing reward information in an action-centric manner (“the value of licking now”) using signals that are synchronized to the lick cycle. Previous studies of MFC reward signaling have inferred value coding upon temporally sustained activity during the period of reward consumption (e.g. Luk and Wallis, 2009). Our findings suggest that neuronal activity in the rostral MFC is temporally sustained during the consumption of rewarding fluids because the animal is licking, and not because of the abstract properties of rewarding fluids.

### Frequency-specific entrainment to licking

Two major rhythms were prevalent in our neuronal recordings. Power was elevated between 2-4 Hz (delta) throughout the performance of the task, but this frequency was not phase locked to the lick cycle (Figure 5). A second major rhythm, which was phase locked to the lick cycle, occurred between 6 and 12 Hz (theta) (Figure 5). The phase of this 6-12 Hz theta rhythm was consistent from lick to lick (Inter-Trial Coherence) and the strength of “phase-locking” was enhanced when rats consumed the higher value reward (Figure 7). Spike activity was coherent to this theta frequency range for all multi-unit recordings that were made simultaneously with the field potential recordings (Figure 8). Theta-range rhythms are a prominent feature of the frontal cortex across species (Cavanagh and Frank, 2014). In rodents, frontal theta signals represent information about behavioral outcomes and performance adjustment (Narayanan et al., 2013; Laubach et al., 2015), interval timing (Parker et al., 2014; Emmons et al., 2016), freezing during fear conditioning (Karalis et al., 2016), and consummatory action (Horst and Laubach, 2013). The results reported here suggest that frontal theta also has a role in reward processing.

### Orolingual aspects of MFC function

The prominence of theta in rodent frontal cortex during consummatory behavior is interesting as rats naturally lick at frequencies between 6 and 8 Hz (Weijnen, 1998) and open/close their jaw in the 5-7 Hz range (Sasamoto et al., 1990). These behaviors are controlled by brainstem central pattern generators (CPGs) (Travers et al., 1997). Other orofacial behaviors such as chewing/mastication (Nakamura and Katakura, 1995), breathing, sniffing and whisking (Moore et al., 2013) also occur in a rhythmic manner and are controlled by CPGs in the brainstem (Moore et al., 2014). These subcortical sensorimotor areas receive projections from the rostral MFC (Yoshida et al., 2009; Haque et al., 2010; Iida et al., 2010) and stimulation of rostral MFC has direct effects on orofacial movement (Brecht et al., 2004; Adachi et al., 2008) and breathing (Hassan et al., 2013). In light of these studies, we propose that one potential function of the rostral MFC might be that it serves as “cingulate motor area” (Shima and Tanji, 1998) controlling orolingual actions and the synchronization between MFC activity and the lick cycle might allow MFC to monitor the consequences of consumption in a temporally precise manner.

### Lick-entrained theta is generated by MFC neurons

Unilateral reversible inactivations decreased theta phase tuning (Figure 10C,E) in the opposite hemisphere, established that these signals depend on neurons within the MFC. Our inactivations were done unilaterally and there were no overt behavioral changes to the animals’ licking behavior during the inactivation sessions. This is in strong contrast to bilateral inactivations of the same cortical area which temporally fragments licking and eliminates the expression of learned incentive contrast effects in the task (Figures 3 and 7 in Parent et al., 2015a). It is not uncommon for unilateral cross-hemispheric inactivations to show less dramatic effects on behavior (Ambroggi et al., 2008), and it was necessary for our interpretations to have the rats maintain their behavior without normal MFC function. Our findings from the inactivation study bolster evidence for a role of MFC in generating theta signals that are synchronized to the lick cycle, in addition to more distal sources such as the olfactory system (Fontainini and Bower, 2006) and hipoocampus (Paz et al. 2008).

## Conclusion

We have shown a role for theta rhythmic activity generated in the rostral part of the rat medial frontal cortex in tracking the act of consumption of liquid sucrose rewards. If these signals encode relative reward value or enable comparisons among different rewards, then they could enable control over food-based decisions and self control over eating. New experiments are needed to investigate these issues. In any case, our findings have clinical implications for diseases associated with MFC dysfunction, e.g. understanding anhedonia in psychiatric diseases such as depression and schizophrenia (Gorwood, 2008) and the loss of control over food intake in obesity (Volkow et al., 2011) and eating disorders such as anorexia (Uher et al., 2004). Our findings also raise an alternative interpretation for studies that have reported reward magnitude coding in the MFC without measuring or accounting for the effects of consummatory behavior on neuronal activity.

## Acknowledgements

We thank Alexxai Kravitz, Catherine Stoodley, and Kyra Swanson for helpful comments on the manuscript.

